# A brain-inspired algorithm enhances automatic speech recognition performance in multi-talker scenes

**DOI:** 10.1101/2025.07.15.664627

**Authors:** Alexander D. Boyd, Kamal Sen

## Abstract

Modern automatic speech recognition (ASR) systems are capable of impressive performance recognizing clean speech but struggle in noisy, multi-talker environments, commonly referred to as the “cocktail party problem.” In contrast, many human listeners can solve this problem, suggesting the existence of a solution in the brain. Here we present a novel approach that uses a brain inspired sound segregation algorithm (BOSSA) as a preprocessing step for a state-of-the-art ASR system (Whisper). We evaluated BOSSA’s impact on ASR accuracy in a spatialized multi-talker scene with one target speaker and two competing maskers, varying the difficulty of the task by changing the target-to-masker ratio. We found that median word error rate improved by up to 54% when the target-to-masker ratio was low. Our results indicate that brain-inspired algorithms have the potential to considerably enhance ASR accuracy in challenging multi-talker scenarios without the need for retraining or fine-tuning existing state-of-the-art ASR systems.

## Main Text

Neuroscience has had a profound influence on the field of artificial intelligence (AI) since its inception (*1, 2*) The most basic elements in artificial neural networks were modeled after neurons in the brain. Feature detection in artificial neural networks drew upon neuroscience discoveries, e.g., orientation selective cells in the visual system. These networks evolved in their computational capabilities by adopting architectures and concepts from neuroscience, e.g., multi-layer “deep” neural networks, learning and memory. Current AI systems are extremely impressive in terms of their performance, outperforming humans in some areas. However, in other areas they have yet to match human-like performance. How and when AI systems will achieve artificial general intelligence remains unclear. A major goal in the emerging field of NeuroAI (*3*) is to advance next generation AI systems with more human-like performance by utilizing insights from neuroscience.

Here, we take a NeuroAI approach towards the cocktail party problem (*4, 5*), i.e., the problem of recognizing a speaker in the midst of competing speakers in a noisy scene. Most people with normal hearing are able to solve this problem with relative ease. In stark contrast, this problem remains extremely challenging for certain populations (e.g., those with hearing loss, ADHD, or autism) as well as for ASR systems (e.g., SIRI or Alexa). Previous studies have employed deep neural networks (DNNs) to address this problem (*6*). However, brain-inspired approaches have the potential to generate much more energy efficient algorithms (*7*). We recently developed a brain inspired sound segregation algorithm (BOSSA) to help solve the cocktail party problem (*8, 9*). BOSSA consists of a multi-stage, experimentally based model of the auditory system, which converts binaural acoustic inputs into neural spike trains. A key feature of this network is a layer of spatially tuned neurons (STNs), which selectively respond to sound from a particular location (azimuth) in space, corresponding to the desired target sound. In the presence of competing sounds from other locations, the response of the STNs at the target location is refined via network inhibition, to generate an approximation of the spike train in response to the “clean” target. Finally, the spike train is reconstructed into an acoustic waveform corresponding to the target sound using signal-processing techniques. This final audio output can be provided to a user/system for speech recognition to improve intelligibility.

We have recently demonstrated that BOSSA leads to robust gains in speech recognition in listeners with and without hearing loss when confronted with challenging multi-talker scenarios (*8, 9*). Here, we evaluate whether BOSSA can also improve the performance of Whisper, a state-of-the-art ASR system (*10*), in multi-talker scenarios.

## Methods

### Stimulus Generation

Stimuli were generated for a given trial by selecting and spatializing three random recordings from the LibriSpeech test-clean audiobook corpus (*11*). Voices and utterances were unique within and across trials. Each recording was spatialized to one of three azimuths (θ ∈ {0°, ±30°}) via convolution with a CIPIC small pinna head-related transfer function sampled from a Knowles Electronics Mannequin for Acoustic Research at 44.1kHz (*12*). Each audio stream was normalized to the same root mean square amplitude and a gain was then applied to the target signal to achieve the desired target-to-masker ratio (TMR). Following TMR adjustment the three audio streams were summed to create a stereo mixture. In the control condition, the stereo mixture was then converted to mono by averaging the left and right channels and downsampled to 16kHz sampling rate to match the input specifications of Whisper. The experimental condition incorporated the additional BOSSA pre-processing step before mono conversion and downsampling. Figure 1 depicts the spatial locations of competing talkers and sequential audio processing steps in presenting stimuli to the Whisper ASR.

**Fig. 1.**
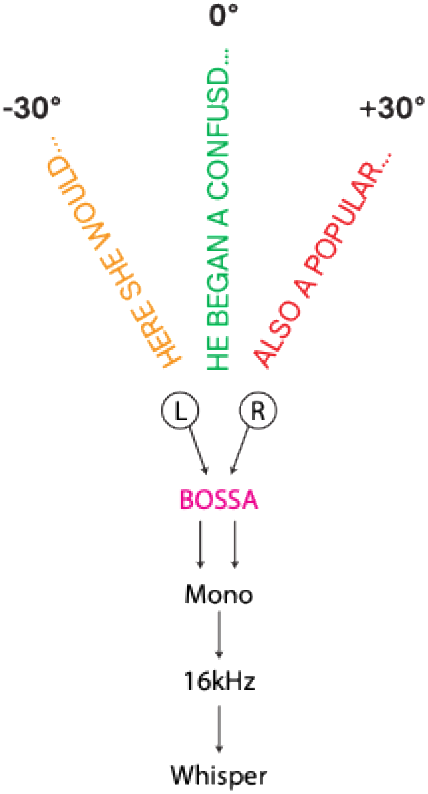
Source configuration and signal flow of three competing talker stimulus. The target source was always located at 0° azimuth. Different colors depict unique voices at each spatial location.

### BOSSA

Drawing on the principles of ideal time-frequency mask estimation, BOSSA processes incoming binaural audio by dividing it into time-frequency bins and selectively applying gain to retain energy from a designated target location while suppressing energy from non-target directions (Figure 2). As outlined in Chou et al. (*8*) the gain for each time-frequency bin is calculated by integrating the predicted activity of STNs, each tuned to a preferred target angle. The real-time implementation of BOSSA used in this experiment included three STNs with preferred source directions of θ ∈ {0°, ±30°}. Another difference from previous versions of BOSSA was the number and range of frequency bands which were reduced to 32 equivalent rectangular bandwidth spaced filters with center frequencies spanning 200 to 8000 Hz. For computational efficiency the real-time version of BOSSA calculated output masks using STN firing rates, instead of the spike trains used in previous versions (*8, 9*).

**Fig. 2.**
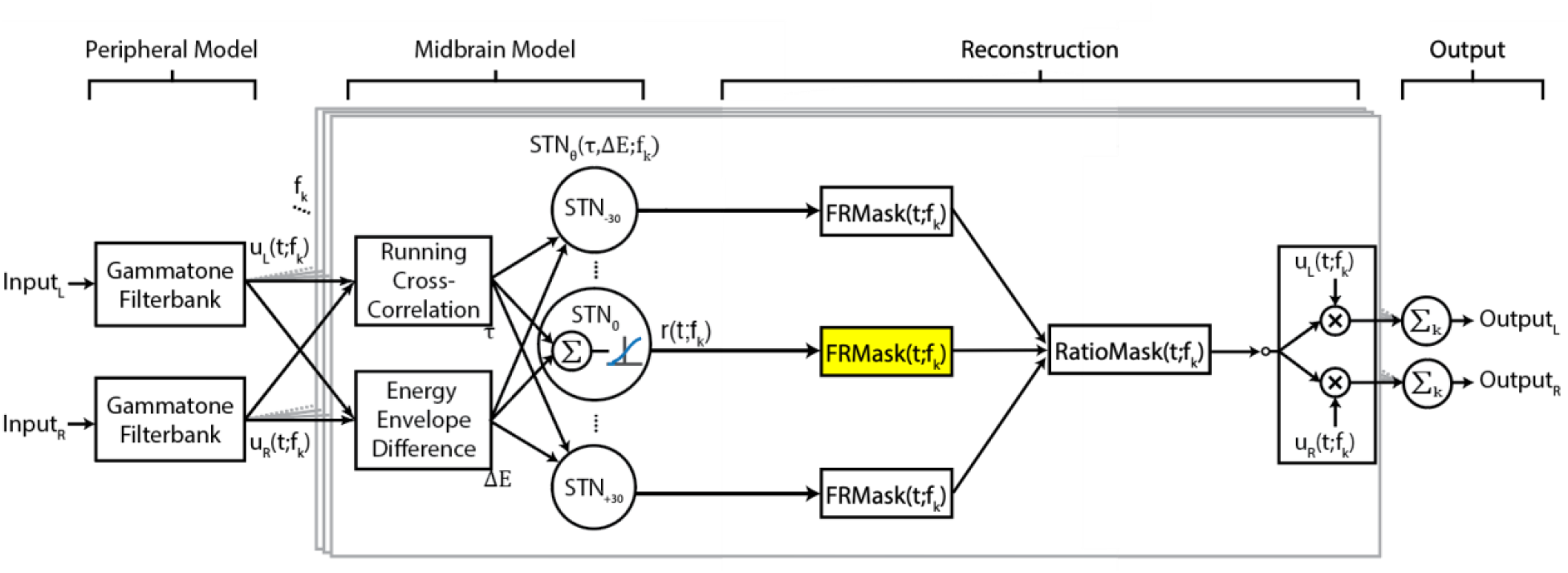
Flow diagram of BOSSA algorithm. Central boxes, outlined in grey, show processing for a single frequency band. The functions *u*_*L*_(*t*; *f*_*k*_) and *u*_*L*_(*t*; *f*_*k*_) are the narrowband signals of the left and right input channels for each frequency channel, and *f*_*k*_denotes the *k*^*th*^ frequency channel. The midbrain model is based on STNs, where each STN has a “best” interaural time and level difference, denoted τ and *ΔE*, respectively. The best interaural values of a neuron depend on the direction *θ* and frequency *f*_*k*_to which the STN is tuned. RatioMask was used for reconstruction. The target STN for reconstruction (yellow highlight) is manually selected as a parameter. The implementation of RatioMask in our analysis involves three sets of STNs, where *θ* ∈ {0, ±30}; other implementations of the model may involve different sets of STNs.

### BOSSA Parameters

The majority of model parameters were fixed to biologically plausible values. Some reconstruction parameters such as internal gains were tuned for Whisper using a bounded genetic algorithm function from the MATLAB Global Optimization Toolbox with “fitness” defined as WER. A time-limited parameter search was conducted at each TMR using a single audio trial per TMR. These stimuli were created with the same procedure described above with the exception that source audiobook recordings were taken from the LibriSpeech dev-clean corpus subset (*11*) instead of test-clean to avoid optimizing for a voice included in testing. We do not claim that this method found the true general optimal parameter set for a given TMR, but rather this process produced generally acceptable solutions while avoiding extensive parameter space explorations.

### Evaluation

The WER was computed for both conditions by comparing the Whisper output transcript against the target talker (0°) reference transcript. Case and punctuation were removed before each WER calculation. WER and word accuracy rate (WAR = 1 -WER) were calculated across 20 trials per TMR as TMR was incremented in 1 dB steps from -5 to +15 dB (equivalent to a signal-to-noise ratio of ∼ -12 to +8 dB). Each trial contained approximately 100 words at an average trial duration of 45.4 seconds. This yielded a total of roughly 2,000 total words tested over the course of ∼15 minutes of speech per TMR. The confidence interval for WER was estimated using non-parametric bootstrap resampling (n=1,000) for both the mean and median WER.

## Results

Figure 3A shows the median WAR across 20 trials as a function of TMR for both the BOSSA-processed and control conditions. These plots show the expected positive correlation between WAR and TMR for both conditions. Additionally, a difference between control and BOSSA-processed audio median WER is seen in TMR conditions from -5 to +5 dB TMR.

**Fig. 3.**
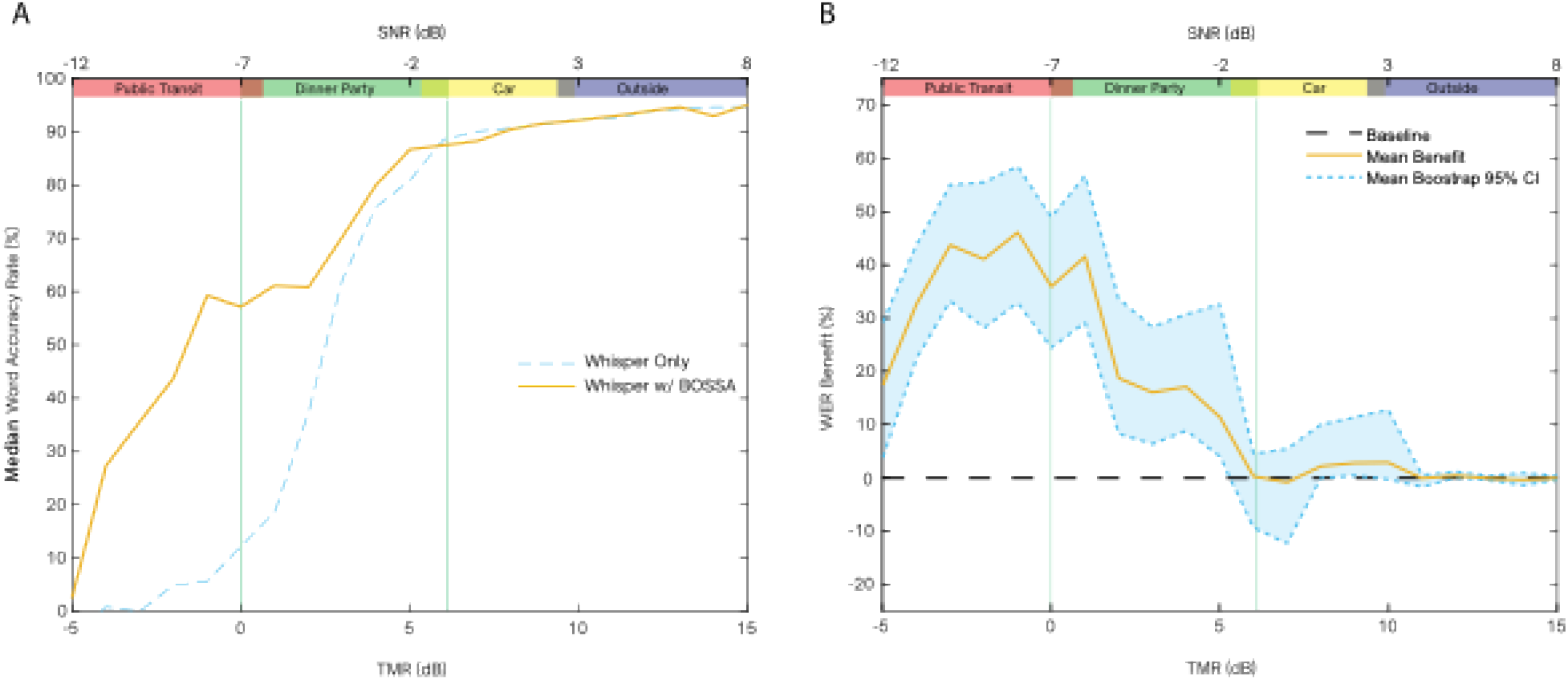
Performance summary of Whisper ASR with and without BOSSA preprocessing. **(A)** Median WAR across 20 trials per TMR with and without BOSSA as TMR increased from -5 to +15 dB. **(B)** Mean WER benefit of BOSSA on ASR performance as a function of TMR with bootstrap 95% confidence intervals. Real-world SNR ranges are displayed above each plot by color bars to provide context.

The effect of BOSSA processing is more clearly displayed in Figure 3B which shows mean WER benefit as a function of TMR. The peak of WER benefit occurs at -1 dB TMR (SNR ∼ - 8dB) where BOSSA processing improved mean and median WER by 46% and 54% respectively. The lower bootstrap confidence interval for mean WER benefit is above 0 from -5 to 5 dB TMR. Additionally, the minimal impact of BOSSA processing for TMRs above 6 dB (SNR > 0) where baseline WAR is already above 85% can be seen.

A more detailed breakdown of WER data and for each TMR can be found in Table S1.

## Discussion

Humans are highly adept at speech recognition even in noisy conditions or extremely challenging conditions involving multiple competing talkers (*8, 9*). ASR systems have long strived to match this impressive capacity (*10, 13*). Although the performance of ASR systems in quiet backgrounds has been impressive for some time, such systems initially appeared to struggle in noisy backgrounds (*14*). Thus, an updated milestone in ASR has been to improve performance in noisy backgrounds to match or exceed human performance. While this goal appeared elusive even a few years ago, recent results indicate that state-of-the-art systems are capable of matching or exceeding human performance in noisy backgrounds in some cases. Specifically, a recent study showed that the state-of-the-art system Whisper outperformed humans in word recognition at an SNR of 0 dB in a background of stationary speech-shaped noise, but not in a more realistic pub noise (*15*). Real-world environments are often characterized by poor SNRs (*16, 17*) and highly variable competing sounds (such as competing speakers). Strategies for improving state of the art ASR in such conditions will further advance the real-world utility of ASR systems.

Here, we adopted a NeuroAI approach to address this problem. NeuroAI approaches have adopted brain-inspired principles to develop and enhance AI systems in a variety of areas (*3*). However, until recently, it had not been applied to the problem of speech recognition. Recently, we developed BOSSA, a brain-inspired algorithm for sound segregation in noisy multi-talker environments, which can improve human speech recognition performance in multi-talker settings with low SNRs (*8, 9*). BOSSA’s ability to improve ASR systems had previously not been evaluated.

Our study found that the integration of BOSSA preprocessing yielded substantial improvements in ASR performance, particularly under conditions characterized by competing speech that was higher in level than that of the target speech. Specifically, in a three-talker scenario the median ASR accuracy improved by up to 54% as a result of BOSSA preprocessing. The TMR range in which BOSSA improved WER (TMR -5 to +5 dB, SNR ∼ -12 to -2 dB) corresponds to real-world noisy settings, such as public transit environments (SNR approximately -10 dB) and bustling food courts (SNR approximately -7.1 dB). These results underline the effectiveness of BOSSA in mitigating the detrimental impact of realistic background noise on speech recognition accuracy.

Moreover, the performance benefits provided by BOSSA come with minimal computational cost. The preprocessing step introduced only a 11.6 millisecond delay per audio frame, (buffer size of 512 samples at 44.1 kHz 16-bit), suggesting that the algorithm can be seamlessly integrated into real-time ASR systems without introducing perceptible delays.

Our findings are significant because they indicate that incorporating a brain-inspired audio preprocessing approach can greatly enhance the robustness of ASR models in cocktail-party scenarios where multiple speakers are present and distinguishing voices is difficult. Importantly, this improvement was achieved without requiring any retraining or fine-tuning of the ASR model itself. This is a major advantage because training state-of-the-art AI can consume enormous amounts of energy. We observed similar improvements when BOSSA preprocessing was applied to other ASRs, e.g., wave2vec (data not shown). Our results show that BOSSA can act as a versatile enhancement to improve speech recognition in noisy environments while maintaining compatibility with existing ASR architectures and minimally affecting performance in less noisy scenarios where accuracy is already high. The fact that audio processing by BOSSA does not assume any specific language also makes it well-suited for ASR systems in other languages, without requiring retraining.

In real-world scenes, multiple sounds sources are usually located in different spatial positions. The success of BOSSA in enhancing the performance of Whisper in noisy backgrounds can be attributed to segregating a target sound from a particular spatial location and providing this spatially segregated output to Whisper. To our knowledge, alternative technologies for spatial segregation, e.g., beamformers and deep neural networks(*6*), have not been evaluated for their effects on Whisper. In our previous study, we showed that the performance of BOSSA, which requires two microphone inputs, provided similar benefits to human listeners as a beamformer with 16 microphones (*8*). Moreover, BOSSA, which is based on a spiking neural network, can be implemented in neuromorphic architectures, which are much more energy efficient compared to deep neural networks (*7*). Thus, BOSSA can provide an energy efficient solution with a compact form factor for enhancing the performance of ASR systems in noisy scenes.

A limitation of the version of BOSSA used in this study is that it used only three STNs to enable real-time application with acceptable latency. Future work should explore software and hardware implementations to increase spatial resolution while maintaining acceptable latency. Another future direction is to add the capability of sound localization. This would allow BOSSA to infer the locations of sound sources in a scene and adaptively allocate the appropriate number of STNs. Finally, it is worth noting that BOSSA did not include standard noise-reduction approaches for background noise, e.g., Wiener filtering or spectral subtraction. Future extensions incorporating such add-ons may further improve performance.

The auditory spatial representation in STNs in BOSSA opens up the ability to incorporate models of auditory spatial attention (*18*) into ASR and, more generally, AI systems. Models of visual attention have been proposed in neuroscience (*19, 20*), and machine learning has adopted the concept of attention, including spatial attention, to dramatically improve the performance of AI systems in computer vision and image processing (*21*). In the visual domain, there is a natural notion of spatial location for an image defined in pixel space, which can be exploited by AI systems. However, in the auditory domain, the spatial location of objects needs to be computed based on acoustic cues. By modeling how the brain computes spatial location from peripheral binaural cues, BOSSA provides a NeuroAI framework for incorporating auditory spatial representations and auditory attention into AI systems. This framework could ultimately be integrated with models of visual attention to develop a supra-modal attention model for audio-visual processing by AI systems.

Overall, our results highlight the potential of adopting a NeuroAI approach to complex auditory scene analysis making ASR and AI systems more applicable to real-world scenes with multiple competing objects, background noise, and clutter.

## Acknowledgments

The authors would like to thank Kenny Chou for his prior work on the BOSSA algorithm. We thank James Galagan and Brian DePasquale for comments on the manuscript.

## Author contributions

Conceptualization: KS, AB

Methodology: KS, AB

Investigation: AB

Visualization: AB

Funding acquisition: KS

Project administration: KS

Supervision: KS

Writing – original draft: KS, AB

Writing – review & editing: KS, AB

## Data and materials availability

The raw data supporting the conclusions of this article will be made available by the authors, without undue reservation.

## Supplementary Materials

**Table S1.**
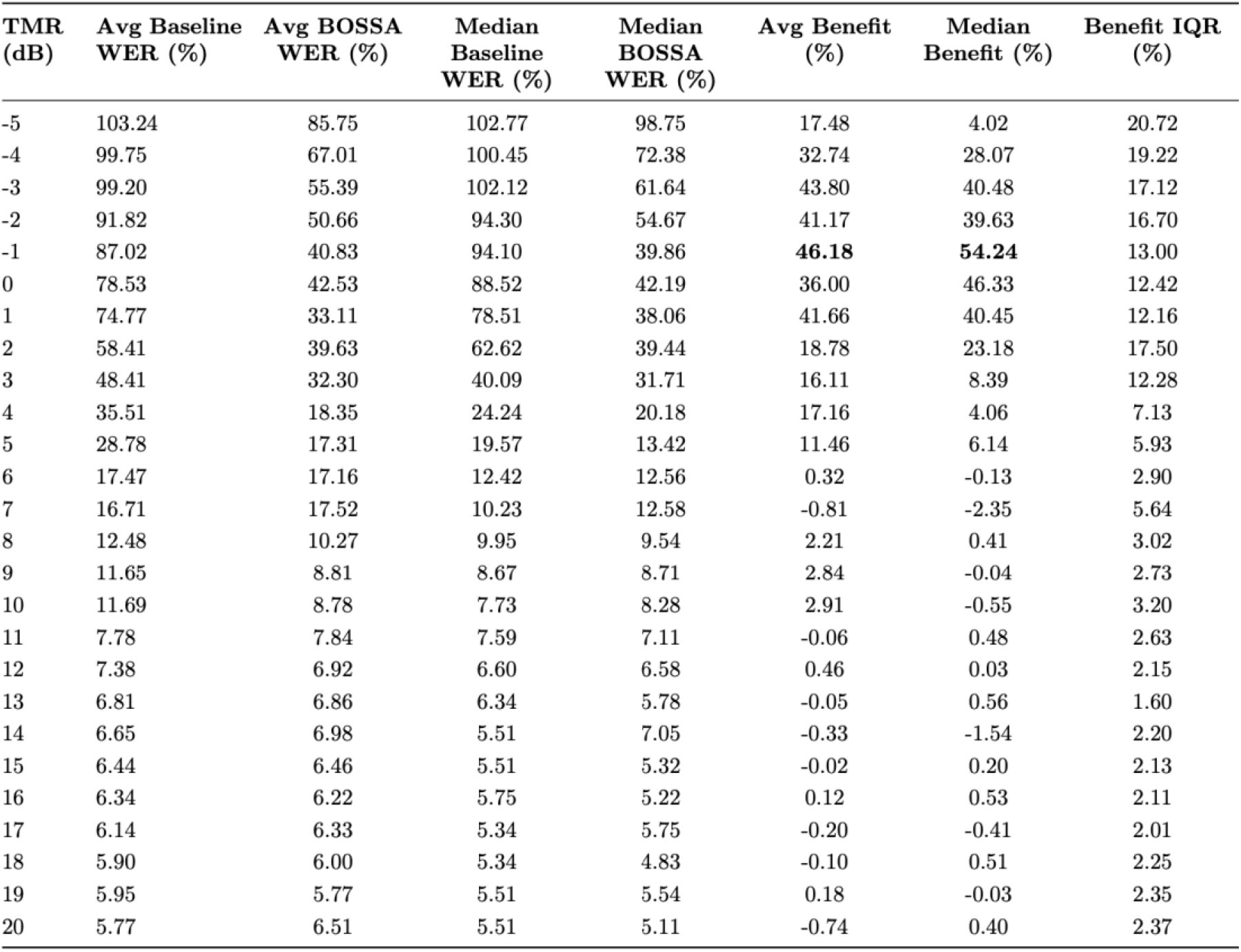
Table summarizing WER results across all tested TMRs. The largest WER benefit values are demarcated in bold.

