## Supplementary Materials Table S1. for "A brain-inspired algorithm enhances automatic speech recognition performance in multi-talker scenes"

### Table S1.

Table summarizing WER results across all tested TMRs. The largest WER benefit values are demarcated in bold.

| TMR (dB) | Avg Baseline WER (%) | Avg BOSSA WER (%) | Median Baseline WER (%) | Median BOSSA WER (%) | Avg Benefit (%) | Median Benefit (%) | Benefit IQR (%) |
|---|---|---|---|---|---|---|---|
| -5 | 103.24 | 85.75 | 102.77 | 98.75 | 17.48 | 4.02 | 20.72 |
| -4 | 99.75 | 67.01 | 100.45 | 72.38 | 32.74 | 28.07 | 19.22 |
| -3 | 99.20 | 55.39 | 102.12 | 61.64 | 43.80 | 40.48 | 17.12 |
| -2 | 91.82 | 50.66 | 94.30 | 54.67 | 41.17 | 39.63 | 16.70 |
| -1 | 87.02 | 40.83 | 94.10 | 39.86 | **46.18** | **54.24** | 13.00 |
| 0 | 78.53 | 42.53 | 88.52 | 42.19 | 36.00 | 46.33 | 12.42 |
| 1 | 74.77 | 33.11 | 78.51 | 38.06 | 41.66 | 40.45 | 12.16 |
| 2 | 58.41 | 39.63 | 62.62 | 39.44 | 18.78 | 23.18 | 17.50 |
| 3 | 48.41 | 32.30 | 40.09 | 31.71 | 16.11 | 8.39 | 12.28 |
| 4 | 35.51 | 18.35 | 24.24 | 20.18 | 17.16 | 4.06 | 7.13 |
| 5 | 28.78 | 17.31 | 19.57 | 13.42 | 11.46 | 6.14 | 5.93 |
| 6 | 17.47 | 17.16 | 12.42 | 12.56 | 0.32 | -0.13 | 2.90 |
| 7 | 16.71 | 17.52 | 10.23 | 12.58 | -0.81 | -2.35 | 5.64 |
| 8 | 12.48 | 10.27 | 9.95 | 9.54 | 2.21 | 0.41 | 3.02 |
| 9 | 11.65 | 8.81 | 8.67 | 8.71 | 2.84 | -0.04 | 2.73 |
| 10 | 11.69 | 8.78 | 7.73 | 8.28 | 2.91 | -0.55 | 3.20 |
| 11 | 7.78 | 7.84 | 7.59 | 7.11 | -0.06 | 0.48 | 2.63 |
| 12 | 7.38 | 6.92 | 6.60 | 6.58 | 0.46 | 0.03 | 2.15 |
| 13 | 6.81 | 6.86 | 6.34 | 5.78 | -0.05 | 0.56 | 1.60 |
| 14 | 6.65 | 6.98 | 5.51 | 7.05 | -0.33 | -1.54 | 2.20 |
| 15 | 6.44 | 6.46 | 5.51 | 5.32 | -0.02 | 0.20 | 2.13 |
| 16 | 6.34 | 6.22 | 5.75 | 5.22 | 0.12 | 0.53 | 2.11 |
| 17 | 6.14 | 6.33 | 5.34 | 5.75 | -0.20 | -0.41 | 2.01 |
| 18 | 5.90 | 6.00 | 5.34 | 4.83 | -0.10 | 0.51 | 2.25 |
| 19 | 5.95 | 5.77 | 5.51 | 5.54 | 0.18 | -0.03 | 2.35 |
| 20 | 5.77 | 6.51 | 5.51 | 5.11 | -0.74 | 0.40 | 2.37 |
